# An explainable unsupervised framework for alignment-free protein classification using sequence embeddings

**DOI:** 10.1101/2022.02.08.478871

**Authors:** Wayland Yeung, Zhongliang Zhou, Liju Mathew, Nathan Gravel, Rahil Taujale, Aarya Venkat, William Lanzilotta, Sheng Li, Natarajan Kannan

## Abstract

Protein classification is a cornerstone of biology that relies heavily on alignment-based comparison of primary sequences. However, the systematic classification of large protein superfamilies is impeded by unique challenges in aligning divergent sequence datasets. We developed an alignment-free approach for sequence analysis and classification using embedding vectors generated from pre-trained protein language models that capture underlying protein structural-functional properties from unsupervised training on millions of biologically-observed sequences. We constructed embedding-based trees (with branch support) which depict hierarchical clustering of protein sequences and infer fast/slow evolving sites through interpretable sequence projections. Applied towards diverse protein superfamilies, embedding tree infers Casein Kinase 1 (CK1) as the basal protein kinase clade, identifies convergent functional motifs shared between divergent phosphatase folds, and infers evolutionary relationships between diverse radical S-Adenosyl-L-Methionine (SAM) enzyme families. Overall results indicate that embedding trees effectively capture global data structures, functioning as a general unsupervised approach for visualizing high-dimensional manifolds.

## MAIN

Alignment-based biological sequence comparison is a foundational aspect of bioinformatics. High-quality sequence alignments are critical for accurate protein classification^1^, function prediction^2^, structure prediction^3^, and evolutionary inference^4^. While alignments excel at comparing closely-related sequences, comparing divergent sequences, especially beyond the “twilight zone” (∼25% sequence identity)^5^ requires sophisticated methods. Profile-based methods such as PSI-BLAST^6^, HMMER^7^, and MMseqs^8^ are capable of comparing sequences within the twilight zone; however, performance depends on alignment parameters such as substitution matrices and gap penalties, derived from prior assumptions about protein evolution. Alignment-free strategies based on word-frequency^9^ or information theory^10^ have been proposed; however, these methods suffer from high false positive rates and cannot capture co-evolutionary information in primary sequences^11^.

Recent advances in representation learning offer a powerful alternative for alignment-free comparison of protein sequences. Using the Transformer neural network architecture^12^, protein language models (LM) such as ESM-1b^13^ and ProtBERT^14^ capture the underlying grammar of biological sequences by training on large, universal proteome databases such as UniProt^15^. These models are trained by masked language modeling in which a random subset of residues in each sequence is replaced with blanks and the model is trained to fill in these blanks using contextual information. During this process, the model translates protein sequences into embedding vectors, which serve as a numerical matrix representation of the original sequence. Sequence embeddings are typically used as input features for machine learning to facilitate supervised predictions of various structure-functional properties^16–18^. Although useful, these methods utilize pre-trained LMs as a black-box feature extractor, resulting in limited interpretability and biological insight. Furthermore, these methods require labeled data, demanding additional labor for curation as well as introducing a potential source of error and bias. Placing an emphasis on unsupervised methods, we developed a set of analytical methods which utilize sequence embeddings as a proxy for protein sequences.

We present a generalized, unsupervised protocol for hierarchical clustering on protein sequence embedding vectors. Benchmark studies across diverse protein superfamilies reveal that sequence embeddings can quantify long-distance evolutionary relationships, beyond the twilight zone of sequence similarity. Visualization of sequence projection vectors reveals cluster-specific sequence motifs, which enable explainability and provide additional support for embedding-based classification. Evaluation of multiple language models reveals that ESM-1b best captures the complexities of protein sequence space. We conclude that embedding-based, alignment-free evolutionary analyses offer a unique set of strengths — well-suited as an orthogonal, complementary approach to traditional alignment-based techniques for protein sequence analysis.

## RESULTS

### Sequence embeddings enable comparisons across long evolutionary distances

We evaluate the ability of protein LMs to model distances between highly divergent protein sequences using the encoder of ESM-1b^13^. LM encoders contain a variable number of Attention blocks^12^ where the majority of interpretable information accumulates at the last Attention block of the encoder **(Figure 1A)**. While previous work has shown that pairwise structural contacts can be inferred as a mathematical function of the attention matrix^19^, we gained additional explainability through sequence projections derived from the embedding vector, calculated downstream to the attention matrix. Sequence projections vectors assign normalized weights to each residue in a given protein sequence **(Supp Method 3**.**1)** which infers important catalytic motifs and fast/slow evolving sites. We later demonstrate this in three diverse protein superfamilies.

**Figure 1.**
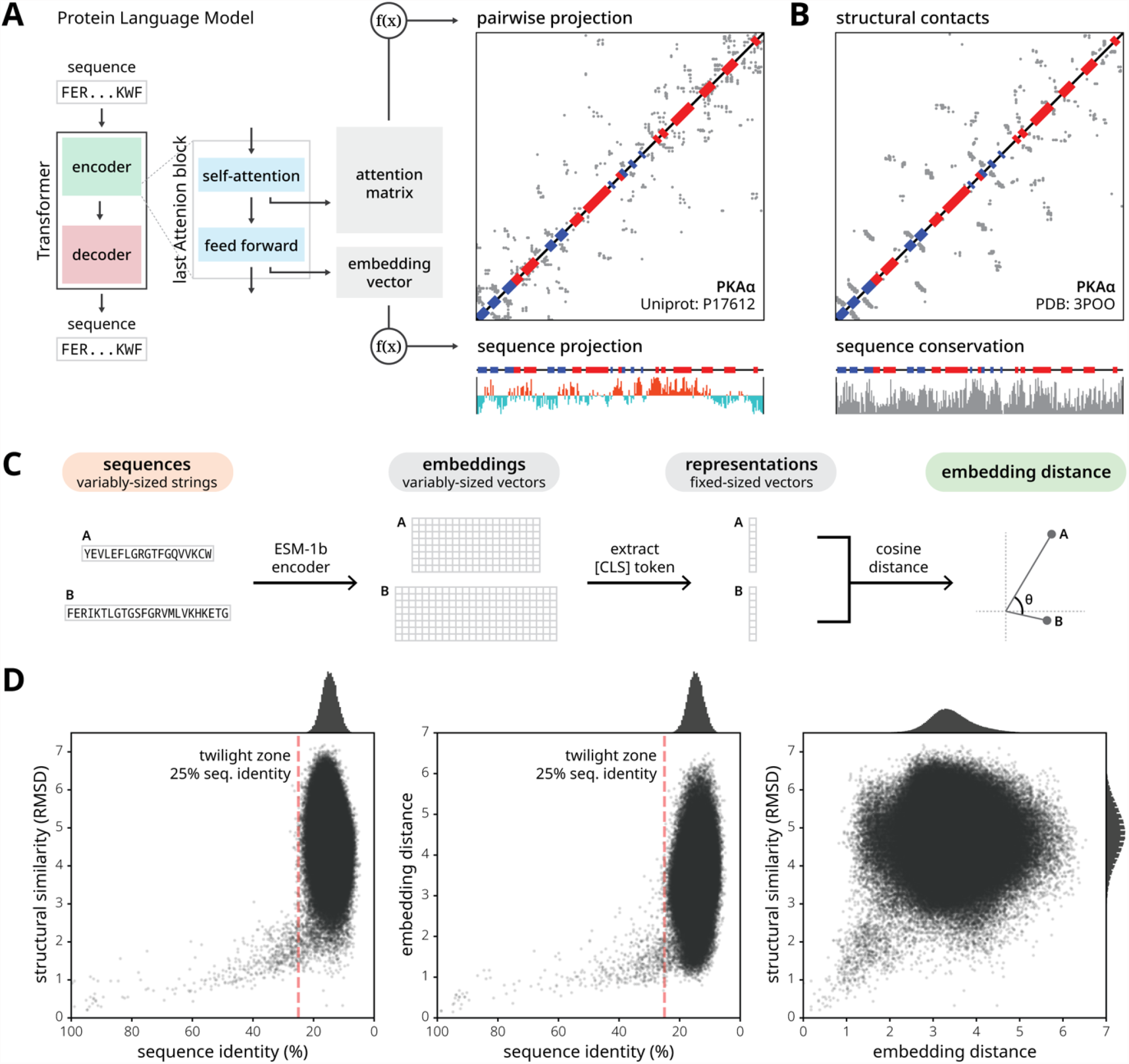
Embedding vectors encode a nuanced description of protein sequence which facilitates comparative analyses between diverse proteins. **(A)** On the far left, we show a graphical representation of a Transformer-based protein Language Model (LM) consisting of an encoder and a decoder module. A zoomed inset depicts major components within the last attention block in the encoder stack. Pairwise contacts can be inferred as a function of the attention matrix^19^. Sequence projections, calculated as a function of the embedding vector, correspond to fast/slow evolving regions. The example was generated from the kinase domain sequence of PKAα (UniProt: P17612) using the ESM-1b model^13^. **(B)** Pairwise structural contacts are shown for the crystal structure of PKAα (PDB: 3POO) defined by C-alpha contacts <7.5 Å. Sequence conservation was calculated by the Jensen-Shannon divergence. **(C)** A graphical overview shows how to calculate embedding distance which provides an embedding-based protocol for pairwise comparison. **(D)** Three scatter plots depict the relationship between sequence and structure (left), sequence and embedding (middle), and embedding and structure (right). Histograms show the distribution of sequence identities, embedding distances, and structural similarities along their respective axes. The twilight zone of sequence identity is marked by a dotted red line at 25% identity^5^. Each point denotes a pairwise comparison between two protein domains. This data was collected by randomly selecting 1,000 proteins from the SCOP (Structural Classification of Proteins) database^20^, provided they were 80-250 residues long with a resolution of 2.3 Å or better.

Embedding vectors facilitate meaningful pairwise comparisons because they encode a nuanced description of protein sequence information. The distance between two embedding vectors can be measured by calculating cosine similarity using the [CLS] special token which is appended before each sequence to capture the sequence-level information during standard preprocessing **(Figure 1C)**. We measured embedding distances between 1000 randomly selected protein domains against standard measures of sequence similarity (percent identity) and structural similarity (RMSD).

As expected, scatterplots show a close relationship between protein sequence and structural similarity that abruptly fades in the twilight zone **(Figure 1D, left)** (sequences below 25% identity)^5,21,22^. Notably, embedding distance is also correlated with sequence identity, displaying a similar boundary at ∼25% **(Figure 1D, middle)**. In contrast, embedding distance and structural similarity (RMSD) display a positive correlation **(Figure 1D, right)**, but instead of a twilight zone, larger variance in embedding distance is observed with increased structural divergence (larger RMSD). This is because sequence embeddings capture a wide range of protein properties beyond 3D structure. Together, these comparisons suggest that protein sequence embeddings can be used as a proxy for sequence and structural similarity metrices and are suitable for comparing sequences in the twilight zone, where traditional alignment-based approaches have proven difficult.

### Unsupervised hierarchical clustering of the protein embedding manifold

Harnessing the unique advantages of protein sequence embeddings, we developed orthogonal methods to facilitate alignment-free evolutionary analyses. We define a hierarchical clustering protocol for constructing embedding trees **(Figure 2, top row)** which provide meaningful organizations of protein sequence datasets and in some instances (see below) can also reflect evolutionary relationships. This protocol has three hyperparameters: the pre-trained LM, representation function, and distance metric **(Supp Methods 2**.**1-2**.**3)**.

**Figure 2.**
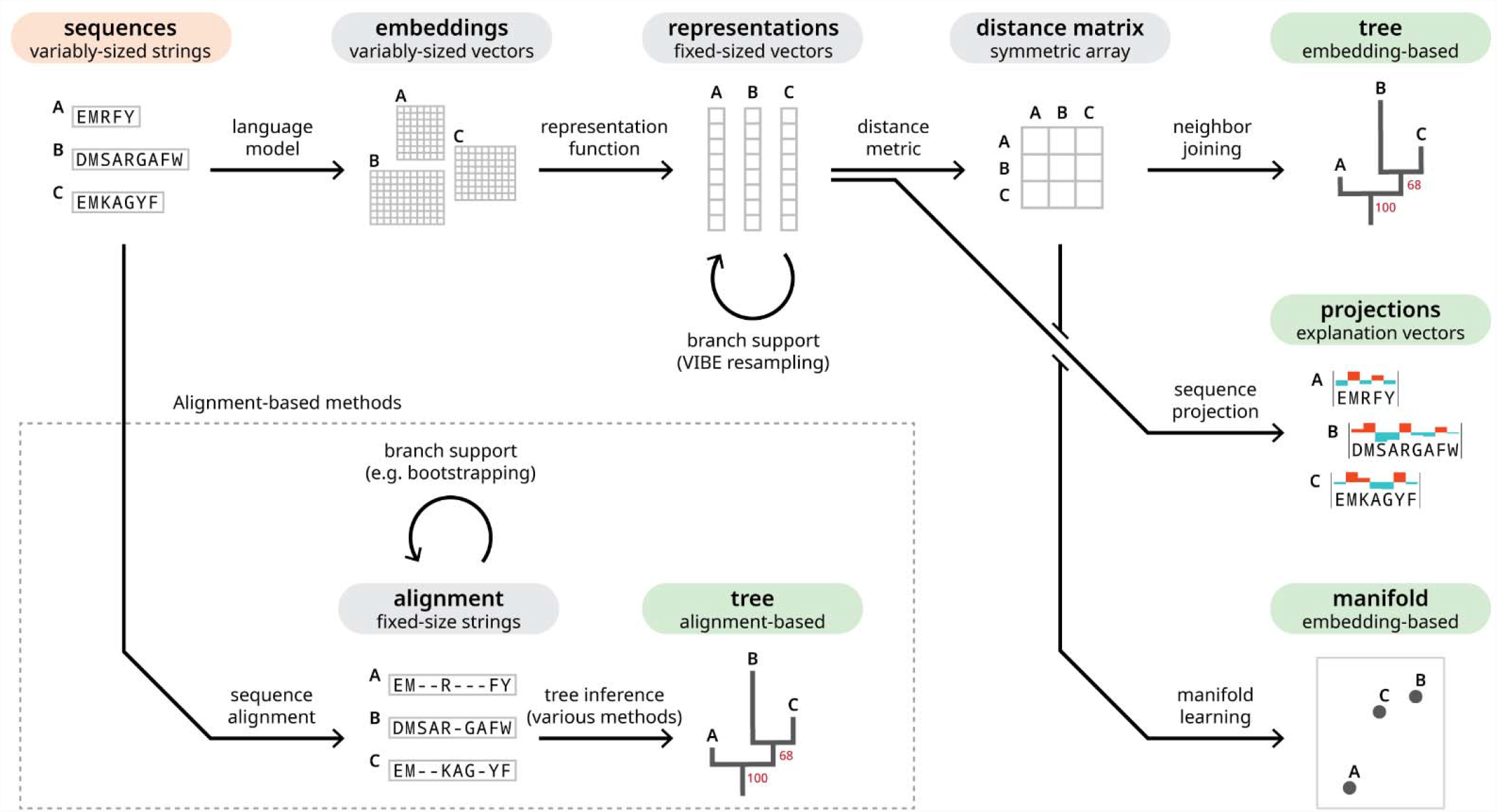
A graphical overview of analysis workflows starting from unaligned protein sequences (label highlighted in red) can lead to four possible endpoints (each label highlighted in green). The top row describes the protocol for creating an embedding tree. Under the representations, the circular arrow denotes our variational autoencoder (VAE)-based strategy for resampling the representation vectors. Resampled representations are used to build replicate trees to calculate branch support, represented by the red number underneath each fork on the tree. Representations, generated from embedding vectors, can also be used to create sequence projections (middle-right) or clustered using manifold learning algorithms such as UMAP (bottom-right). The branching route in the bottom row depicts a more traditional protocol for creating trees using multiple sequence alignments. There are many diverse algorithms for inferring trees using sequence alignments^4^. There are also various methods for resampling data to build replicate trees (such as bootstrapping) which is required for branch support calculations^23^.

We systematically assessed all hyperparameter combinations across diverse case studies using three enzyme superfamilies **(Supp Table S1-S6)**, individually discussed over the next three sections. We used a variety of quantitative measures such as Sackin’s index^24^, treeness^25^, and silhouette coefficient^26^ to evaluate embedding trees **(Supp Methods 3**.**3-3**.**7)**, which are also a general strategy for visualizing high-dimensional datasets — representing pairwise relationships using cophenetic distance. To measure how well the tree preserves all pairwise distances observed in the original data, we quantify the Pearson’s correlation of the tree’s distance matrix versus the representation’s distance matrix. Using the same method, we also compared against manifold learning algorithms such as UMAP (Uniform Manifold Approximation and Projection)^27,28^.

Across all sequence datasets, the ESM-1b model consistently produced trees that agree with previously established protein classifications schemes based on silhouette coefficient **(Supp File S1-S8)**, while also proposing new relationships. Although some LMs such as ProtBERT can be fine-tuned to gain better performance for specific tasks **(Supp Methods 2**.**5)**, fine-tuned LMs did not yield significant improvements in embedding trees **(Supp Table S7-8; Supp File S9-10)**. Given the overall performance of ESM-1b, all analyses throughout this study utilized this LM. Meaningfully compressing embedding vectors^29^ and defining a unified distance metric^30^ are both non-trivial problems. Consequently, the optimal representation function and distance metric varied across different protein datasets.

Upon identifying an optimal tree, we quantify clustering confidence using a variational autoencoder (VAE)-based strategy. The confidence of each split is measured using VIBE (VAE-Implemented Branch support Estimation) **(Figure 2, top-middle)** — conceptually similar to bootstrap support, used in alignment-based phylogenies. As a generative model, the VAE learns the latent distribution of a given set of representations^31^, then resamples the distribution to generate replicate trees. We assign a value to each branch of the original tree, indicating the percentage of replicate trees which also exhibited the same corresponding bipartition. This is a particularly stringent metric which does not consider similar bipartitions if an exact match is not present.

### Embedding trees infer the earliest diverging protein kinase group

We applied our methods towards the protein kinase superfamily — an important gene family which plays diverse roles in cellular signaling and disease. Most protein kinases are classified into nine major groups based on sequence similarity^33^. Outside the protein kinase superfamily, lipid and small molecule kinases are distant relatives which conserve a similar bilobal structure^34^. Although structure-functional similarities strongly imply evolutionary relationships between all kinase-fold enzymes, further characterization has eluded traditional phylogenetic methods.

We built an embedding tree of ∼550 human kinase-fold enzymes using unaligned protein sequences, trimmed to the conserved catalytic domain. The optimal tree organizes sequences into nine major groups **(Figure 3A)** where the inferred between-group relationships are largely consistent with the widely-accepted alignment-based phylogeny^33^. In comparison, untrimmed sequences yield a trivial topology **(Supp File S2; Supp Table S2)**, as meaningful evolutionary analyses require a common frame of reference. To further evaluate confidence, we generated 500 replicates for the kinase domain tree. Sequences from the “Others” category (not belonging to the major protein kinase groups) showed unstable placement across replicates. These rogue taxa are known to decrease branch support^35^, thus we used two common strategies to resolve this issue. The first strategy was to prune rogues from all trees prior to calculating VIBEs, while the second was to rebuild the tree excluding rogue sequences from the dataset **(Figure 3B)**. For both trees, at least 60% of replicates place RGC, TKL, and TK into a monophyletic clade, also placing CAMK and AGC as sister clades — consistent with the existing phylogeny^33^. Extending beyond the existing model, we included an evolutionary outgroup of lipid and small molecule kinases. The placement of CK1 kinases in both topologies infer that CK1 is the earliest diverging protein kinase group, which is further supported by CK1-specific divergence in the substrate binding lobe^36^ and its apparent substrate promiscuity and constitutive activity^37^.

**Figure 3.**
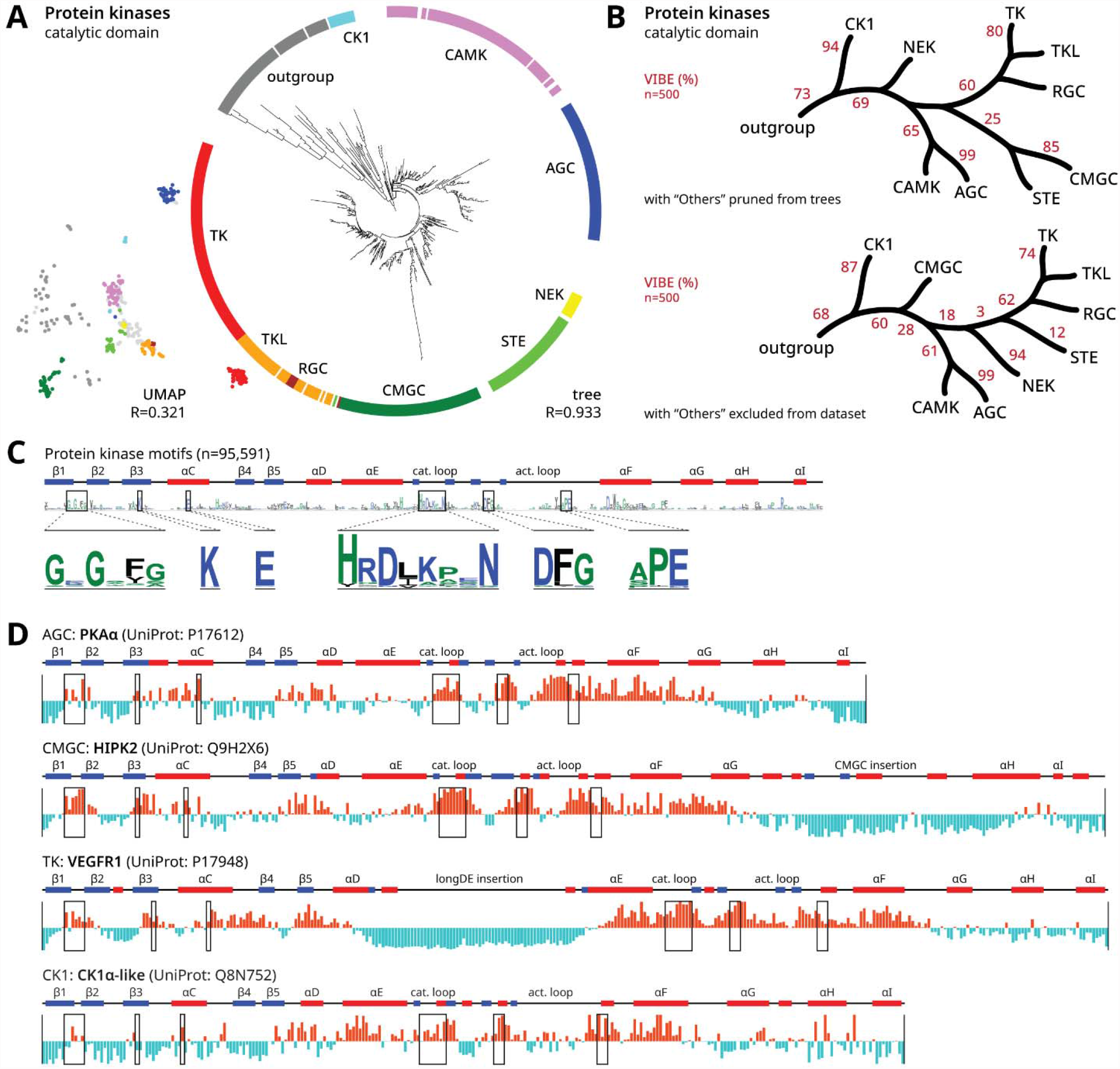
Embedding-based analysis of the human protein kinases. **(A)** An embedding tree of the human protein kinase domains in a circular layout with major groups labeled. This tree was generated using the sum_spec representation function and TS-SS (triangle similarity sector similarity) distance^32^. To the left of the tree, we plot a UMAP projection using the same dataset. At the bottom of each graph, we provide the correlation coefficient which quantifies how well the all-vs-all pairwise distances denoted by each visualization reflects the pairwise distances from the original dataset. The full tree is provided in **Supp File S1** and full technical details are provided in **Supp Table S1. (B)** Stylized trees showing the major kinase groups with VIBEs indicated by the red percentage values. The top topology was inferred by pruning unclassified kinases (“Others” group); full tree is provided in **Supp File S3**. The bottom topology was inferred by excluding the unclassified from the sequence set; full tree is provided in **Supp File S4** and full technical details are provided in **Supp Table S3. (C)** Six major protein kinase motifs are shown within zoomed insets of a sequence logo, generated from an alignment of 95,591 kinase sequences. Above the logo plot, we show secondary structure elements across the kinase domain with α-helices (red), β-sheets (blue), and loops (black). **(D)** Sequence projections for three diverse protein kinase sequences: PKAα, HIPK2, VEGFR1, and CK1α-like. Positive peaks are shown in bright red and negative peaks are shown in icy blue. Sequence regions corresponding to the six major protein kinase motifs are designated by boxes. We note that CK1 kinases lack the APE motif and instead conserve a CK1-specific SIN motif at the equivalent position. Based on the optimal tree parameters, sequence projections were calculated using the sum_spec representation function.

Relationships between sequence embeddings can also be visualized by manifold learning algorithms such as UMAP^27^. We compare against our tree-based method by creating a UMAP projection from the same dataset. The tree-based layout is superior at preserving pairwise distance information, facilitating a more accurate depiction of the underlying manifold. In the kinase domain dataset, all-vs-all pairwise distances from the UMAP projection are weakly correlated to the original data, quantified by a Pearson’s correlation coefficient of 0.366, compared to 0.926 for the tree **(Figure 3A)**. While pairwise distances in UMAP scatterplots are represented by Euclidean distance, pairwise distances in circular trees are represented by cophenetic distance, the sum of branch lengths along the shortest path between two points. Branch length is solely represented by distance across the radial axis, while the circular axis and number of edges do not matter^38^.

Sequence projections provide further explainability for embedding-based analyses. The sequence projection quantifies how strongly a given representation vector weights each residue of a protein sequence. Weights are correlated to fast/slow evolving sites. Most kinases share a common set of sequence motifs such as the nucleotide-binding G-loop motif and catalytic motifs **(Figure 3C)**^39^. A projection of archetypical kinase PKA-α reveals positive peaks for kinase-conserved motifs **(Figure 3D)**. We observe similar peaks for HIPK2 which has a CMGC-specific insertion region towards the C-terminal of the kinase domain^40^ and VEGFR1 which has the longDE insertion towards the center of the kinase domain^41^. These fast-evolving insertion regions correspond to negative peaks. A sequence projection of CK1α-like kinase also highlights protein kinase motifs, albeit with its own unique variations. While determining fast/slow evolving sites typically require a sequence alignment, protein LMs delineate this information without an alignment, functioning as unsupervised learners for fast/slow evolving sites.

### Embedding trees capture similarities between protein folds in protein phosphatases

To further demonstrate the applicability of embedding trees, we generated trees for phosphatase enzymes, which, unlike kinases, adopt distinct structural folds^42^. Out of ∼200 human phosphatases, roughly half adopt the CC1 fold, while only one adopts the RTR1 or PHP fold. The salient heterogeneity of structural folds suggests that phosphatases emerged independently multiple times throughout evolution.

We constructed an embedding tree spanning all ten structural folds using the catalytic domain sequences (Figure 4A). VIBEs were calculated using a filtered dataset which excludes two rogue taxa (Figure 4B). The grouping of the three cysteine-based phosphatase folds (CC1, CC2, and CC3) was supported by 95% of replicates. Within all three structural folds, sequence projections revealed a positive peak at the shared CxxxxxR catalytic motif (Figure 4C). Catalytic similarities between CC1, CC2, and CC3 phosphatases likely arose via convergent evolution — CC2 is more structurally related to bacterial arsenate reductases, while CC3 emerged from bacterial rhodanese-like enzymes^43^. At the opposite end of the tree, PPPL, PPM, and AP were placed into a distinct cluster supported by 66% of replicates. PPPL and PPM phosphatases are phosphoserine/threonine-specific^44^, while AP phosphatases act on phosphotyrosine^45^ with possible phosphoserine/threonine activity based on substrate binding specificities^46^. While these enzymes share similar substrates, the embedding-based similarities between these three folds are not immediately obvious.

**Figure 4.**
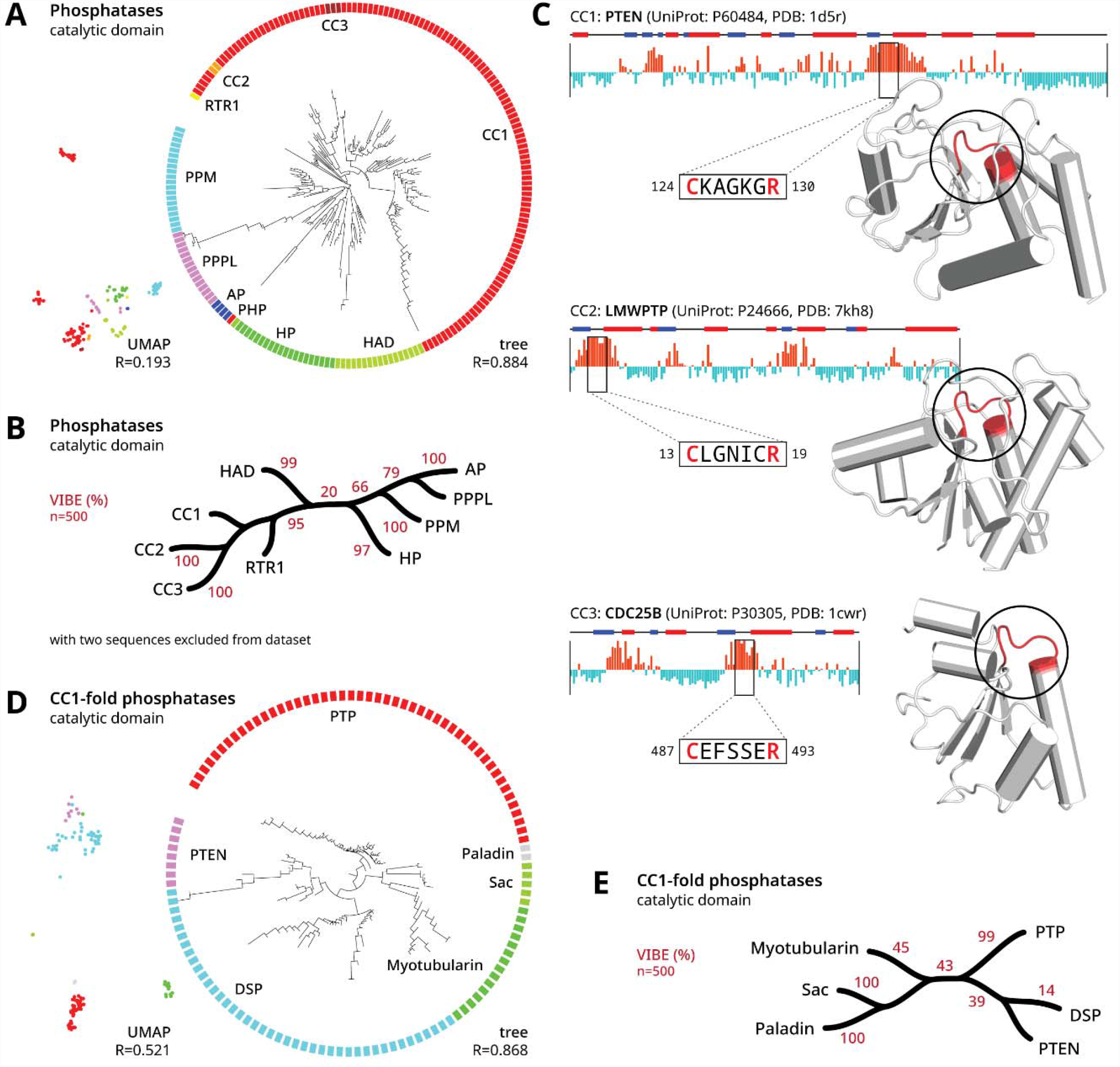
Embedding-based analysis of the human phosphatases. **(A)** An embedding tree of the human phosphatases in circular layout, generated from all human phosphatases enzymes spanning ten structural folds. This tree was generated using the avg_seq representation function and cosine distance. To the left of the tree, we plot a UMAP projection using the same dataset. The full tree is provided in Supp File S5 and full technical details are provided in Supp Table S4. **(B)** A stylized tree showing the phosphatase folds with VIBEs indicated by the red percentage values. This topology was inferred by excluding two rogue taxa, O60729 and Q9NRX4, from the sequence set. The full tree is provided in Supp File S6. **(C)** Sequence projections for a representative CC1, CC2, and CC3-fold phosphatase. The conserved CxxxxxR motif corresponds to a positive peak in all three enzymes, despite adopting different protein folds. Crystal structures of each enzyme are shown with the CxxxxxR motif circled and highlighted red. Based on the optimal tree parameters, sequence projections were calculated using the avg_seq representation function. **(D)** An embedding tree and UMAP projection of the human CC1-fold phosphatases. This tree was generated using the [CLF] representation function and TS-SS distance. The full tree is provided in Supp File S7 and full technical details are provided in Supp Table S5. **(E)** A stylized tree showing the CC1 phosphatase families with VIBEs indicated by the red percentage values.

We also constructed another embedding tree from a reduced dataset only containing CC1-fold phosphatases which adopt a conserved structural fold, implying a common evolutionary origin. Consistent with an alignment-based phylogeny^42^, the embedding tree identified five major clades across the six families. Notably, DSP is a paraphyletic group and shares a clade with PTEN (Figure 4D). VIBEs of the five major clades ranged from 39-100% (Figure 4E). While the CC1-fold tree showed evolutionary relationships, embedding trees for highly divergent sequence sets should be interpreted with caution as similarities can arise from alternative sources such as convergent evolution. Even if evolutionary inferences cannot be made, results can still be interpreted as hierarchical clustering.

### An initial characterization of the radical SAM enzyme superfamily

We apply embedding-based methods towards an initial evolutionary characterization of the radical S-Adenosyl-L-Methionine (SAM) enzyme superfamily. SAM enzymes are present in all domains of life, catalyzing radical chemistry towards a wide variety of essential biological functions^47^. The catalytic core domain of radical SAM enzymes adopts a TIM barrel (α/β barrel) fold with varying numbers of α/β pairs, and a conserved iron-sulfur cluster binding motif, CxxxCxΦC, where Φ denotes an aromatic residue^48^. Family-specific insertions and deletions add additional structural variance, making a superfamily-scale alignment difficult. We curated a dataset of diverse radical SAM enzymes using available protein structures and the AlphaFold2 database^49^. To establish a common frame of reference, we trimmed each sequence to the core catalytic domain, removing any domain extensions or accessory domains.

Despite only utilizing the core domain, an embedding tree of the radical SAM superfamily organized enzymes into structure-functionally similar groups (Figure 5A) with good VIBEs (Figure 5B). For instance, families which specialize in methyl or sulfur transfer (B12-binding, MTaseA, LipA, and MTTase families)^48^ were placed in a single clade. Placed in the neighboring clade, some HemN enzymes also catalyze methyl transfer^50,51^. The HemN and Elp families have reported sequence similarity^52^, while Elp and BAT families both conserve extended TIM barrel folds. Additionally, many Elp and BATS enzymes contain alterations to the canonical CxxxCxΦC motif^53^. Viperin and SPASM families both conserve a C-terminal extension which facilitates family-specific functionalities^54^. Viperin is placed closest to the MoaA subfamily (within the SPASM family); both of which act on nucleotide substrates^55^. Activating enzyme and QueE family members sometimes adopt a “Tiny TIM’’ minimal core fold^56^. QueE and TYW1 families are also closely grouped together; both families are involved in tRNA biosynthesis and hypermodification^57^.

**Figure 5.**
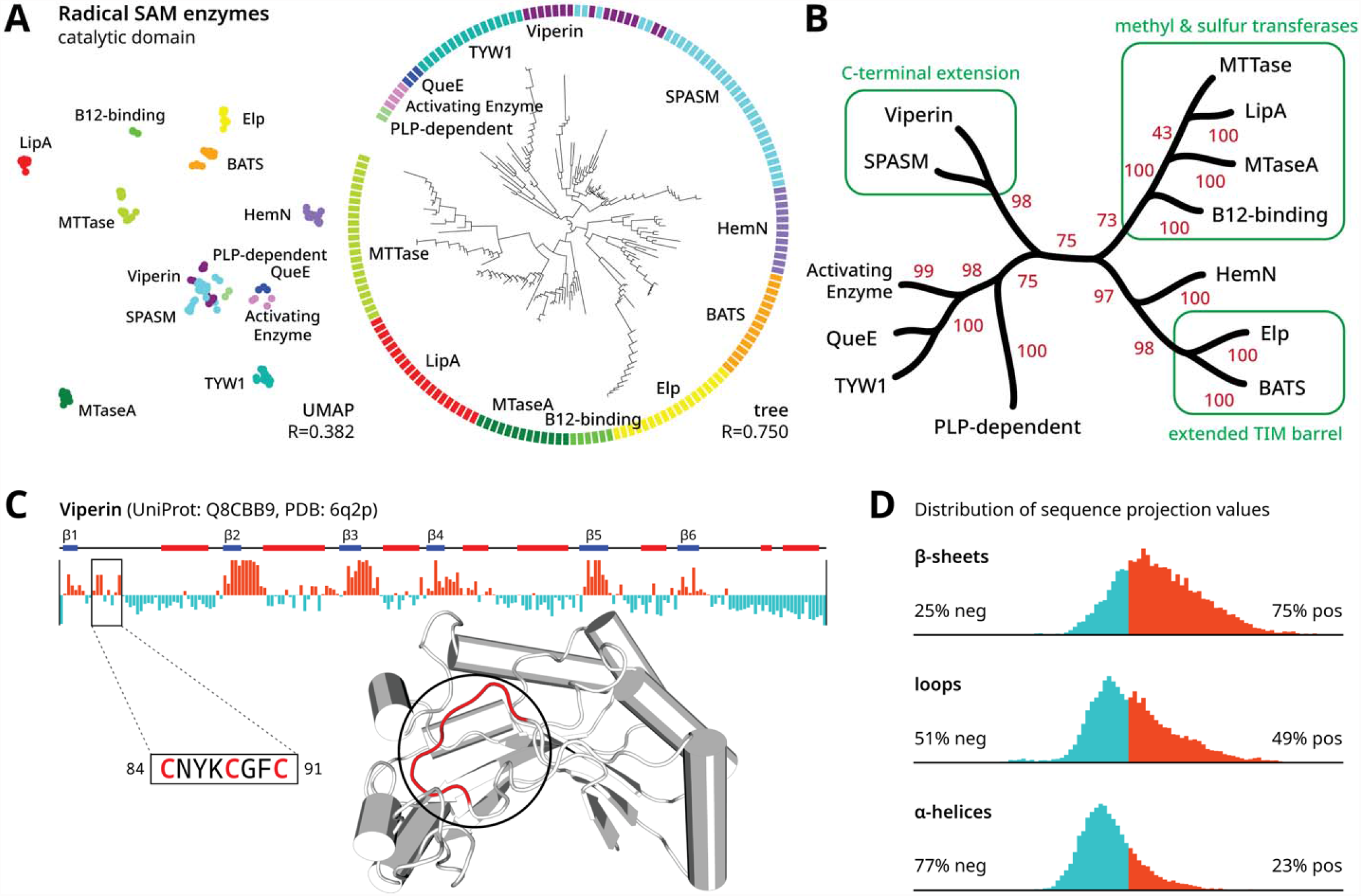
Embedding-based analysis of the diverse radical SAM enzymes. **(A)** An embedding tree of radical SAM enzymes in a circular layout with major groups labeled. This tree was generated using the [CLF] representation function and cosine distance. To the left of the tree, we plot a UMAP projection using the same dataset. At the bottom of each graph, we provide the correlation coefficient which quantifies how well the all-vs-all pairwise distances denoted by each visualization reflect the pairwise distances from the underlying dataset. The full tree is provided in Supp File S8 and full technical details are provided in Supp Table S6. **(B)** A condensed tree showing various families of radical SAM enzymes. VIBEs are indicated by the red percentage values. Select structure-functional annotations are shown in green. **(C)** A sequence projection for a representative radical SAM enzyme. The conserved iron-sulfur cluster binding motif, CxxxCxΦC, corresponds to a positive peak, designated by the zoomed inset. A crystal structure of mouse Viperin is shown with the CxxxCxΦC motif circled and highlighted red. Based on the optimal tree parameters, sequence projections were calculated using the [CLF] representation function. **(D)** We plot histograms of sequence projection values across our dataset of diverse radical SAM enzymes, stratified by the secondary structure at each residue.

Sequence projections across diverse radical SAM enzymes place a positive peak at the conserved CxxxCxΦC motif (Figure 5C). Positive peaks also tend to fall on β-sheets (Figure 5D) extending onto each proceeding loop. These regions correspond to previously identified SAM binding sites found in all radical SAM enzymes, as well as family-specific motifs which facilitate unique family-specific chemistry^58,59^. This trend suggests that the usage of β-sheets in substrate binding and catalysis may be a shared feature across the radical SAM superfamily. Although β-sheets are more conserved than loops and helices^60,61^, sequence projections on other globular protein superfamilies show comparatively weaker association with β-sheets (Supp File S11).

## DISCUSSION

We present an arsenal of orthogonal techniques, listed in Table 1, for alignment-free protein sequence analysis by utilizing sequence embeddings as a proxy for actual amino acid sequences. Throughout diverse case studies, embedding vectors appear most suited for modeling long-distance evolutionary relationships (Figure 1D), allowing us to infer a single tree containing all human kinase-fold enzymes (Figure 3A), identify similarities between divergent phosphatases structural folds which likely arise by convergent evolution (Figure 4A-C), and infer the initial tree of the radical SAM enzyme superfamily (Figure 5A-B). Across all case studies, closely-related proteins had a tendency towards unbalanced, ladder-like topologies with zero branch length tips, suggesting that embedding trees do not have the capacity to resolve closely-related proteins within the same family. While our analyses only utilized shared catalytic domains, a focused analysis on closely-related sequences may benefit from embeddings that include shared regions beyond the catalytic domain. In comparison, while sequence alignments-based approaches are not well suited for long-range evolutionary inference, they work well on closely related sequences. A combination of alignment and alignment-free embedding approaches are expected to advance the frontiers of sequence analysis.

**Table 1.** Comparison of equivalent methods for protein sequence analysis. For a diverse range of sequence alignment-based methods, we define an analogous embedding-based approach.

|  | alignment-based methods | embedding-based methods |
| --- | --- | --- |
| sequence comparison | sequence alignment<sup>62</sup> | embedding distance (Supp Method 2.3) |
| residue conservation | statistical entropy (e.g. Shannon entropy, Kullback-Leibler divergence, Jensen-Shannon divergence)<sup>2</sup><br>sequence logo<sup>63</sup> | sequence projections (Supp Method 3.1) |
| sequence clustering | sequence similarity networks<sup>64</sup> | embedding trees (Supp Method 2.4)<br>manifold learning (e.g. t-SNE<sup>65</sup>, UMAP<sup>27,28</sup>) (Supp Method 3.2) |
| tree inference | probabilistic methods (e.g. maximum likelihood<sup>66</sup>, Bayesian inference<sup>67</sup>)<br>distance matrix methods (e.g. neighbor-joining<sup>68</sup>) | embedding trees (assuming sufficient orthogonal evidence) (Supp Method 2.0) |
| branch support | bootstrapping<sup>69</sup> | VIBEs (Supp Method 3.7) |

Embedding trees indicate that the protein LM provides a reasonable model of the theoretical evolutionary landscape. Although it is technically possible to build embedding trees from any sequence dataset, evolutionary inference should only be invoked if common ancestry can be supported by orthogonal evidence. Common ancestry between kinase fold enzymes is supported by a highly conserved structural fold and sequence motifs^34^. Although phosphatase enzymes share a common catalytic function, different folds utilize different mechanisms which indicate that these enzymes independently emerged multiple times throughout evolution^42^. Consequently, the phosphatase fold tree only should be interpreted as clustering. Despite methodological differences, many established principles in phylogenetic analyses remain relevant such as generating replicate trees and being vigilant towards confounds such as long-branch attraction^70^.

Beyond biological applications, our study provides useful methods for explainable machine learning. Tree-based visualizations more accurately capture the global data structure of high-dimensional data compared to manifold learning algorithms such as UMAP. Citing a major difference, our tree-based method does not frame manifold visualization as a dimensionality reduction problem; trees are inherently capable of depicting high dimensional relationships without assuming an underlying geometry. Sequence projections can also be used as explainability vectors. Not requiring backward gradient calculations, our method demonstrates superior computational efficiency and simplicity. By showcasing these new applications, we hope to promote the development of better LMs. Recent results have proposed mechanisms for generating fixed-sized embeddings from variable-size inputs^71,72^ which would potentially exclude the need for representation functions. Further advances in the field of representation learning are expected to improve the unsupervised classification of large protein families.

## METHODS

### Data collection and preprocessing

The sequence dataset of 558 human kinase-fold enzymes^73^ and 204 human phosphatase sequences^42^ were derived from previously published studies. Our dataset of 179 taxonomically diverse radical SAM enzymes was manually curated based on a previous sequence clustering study^47^. Core domain segments were manually identified and trimmed based on all available crystal structures and AlphaFold2^49^ models. Secondary structure annotations were assigned based on AlphaFold2 models using the DSSP algorithm^74^. Further details about data curation are listed in Supp Methods 1.0-1.4.

### Calculating embedding trees

Following sequence dataset curation, the sequences were converted into embedding vectors using a Transformer-based protein LM (Supp Methods 2.1). Specifically, the embedding vector is the final hidden state generated from the last layer of the encoder module.

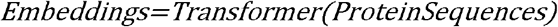

Each embedding is a two-dimensional matrix. The token dimension encodes one token for each residue of the original sequence plus additional special tokens which are appended during preprocessing, while the embedding dimension encodes information about each token. The number of special tokens and the size of the embedding dimension will vary depending on the specific LM used. To enable direct comparisons between embeddings, we derive representation functions to summarize the information encoded within the variably-sized embedding vectors into fixed-sized representation vectors. This is conceptually similar to pooling operations, typically used to condense information within convolutional architectures. Each representation function is applied along the token dimension of the embedding, defined as a function of the special tokens or sequence tokens (Supp Methods 2.2). We explored 8 pretrained protein LMs and 9 different representation functions. After sampling all compatible pairs of protein LM and representation function, we generated 56 unique sets of representation vectors for each sequence dataset.

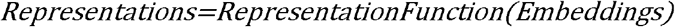

We explored a variety of distance metrics to calculate an all-vs-all distance matrix from the representation vectors: Euclidean distance, cosine distance, Manhattan distance, geodesic distance, and TS-SS^32^. Details pertaining to each distance metric are provided in (Supp Methods 2.3). We sampled each unique combination of representation vectors and distance metrics to generate 280 unique distance matrices for each sequence dataset.

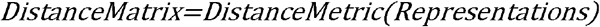

Trees were generated from distance matrices using the neighbor-joining algorithm^68^.

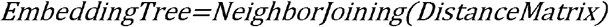

### Evaluating branch support

The statistical confidence of a given bipartition of a tree can be evaluated using a VAE. We trained a VAE on a fixed set of representation vectors to resample replicate representation vectors. By learning a smooth latent state representation from the input data, the VAE becomes capable of regenerating the input data using the reparameterization trick which allows backpropagation through a random node. This unique property of VAE enabled us to resample the original input with any desired number of replicates. To accurately model the underlying space of the protein representations, the VAE is trained on optimizing a combination of Mean Square Error (MSE), Kullback-Leibler divergence (KLD), and TS-SS Error (TSE). We applied the cosine annealing^75^ to control for the weight of the KLD loss term. The detailed model structure can be found in (Supp Methods 3.7).

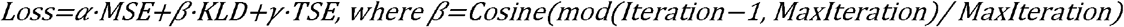

We trained a separate VAE for each unique dataset of representations. VAEs were trained for 20,000 epochs with early stopping patience of 1000 epochs. Resampled representation vectors generated from the final model were used to build 500 replicate trees. Branch support values were assigned to the original tree using the replicate trees. We refer to this procedure and confidence metric as VAE Implemented Branch Support Estimation (VIBE).

### Calculating sequence projections

To understand how a representation vector (generated from a given representation function) encodes an embedding, we calculated the cosine distance between the representation vector and each sequence token of the embedding (Supp Methods 3.1). The resulting sequence projection vector has the same size as the protein sequence corresponding to the embedding. Sequence projections were further standardized to facilitate comparisons between sequences.

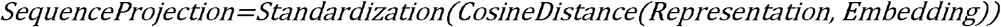

## ACKNOWLEDGEMENTS

This research was supported by ARO funding to SL (W911NF-21-1-0028) and NIH funding to WL (R01 GM124203) and NK (R35 GM139656).

We thank Dr. Liang Liu for valuable feedback and discussions.

## AUTHOR INFORMATION

### Contributions

WY, ZZ, SL and NK conceived the project. WY and ZZ implemented the algorithms and methods. WY, ZZ, LM, NG, RT, and AA curated sequence datasets and analyzed results. WY drafted the manuscript with edits from ZZ, LM, SL, and NK. NK, SL, and WL provided funding. SL and NK provided project supervision. All authors read and approved the manuscript.

## Notes

### Competing Interest Statement

The authors have declared no competing interest.

